# Emotional context sculpts action goal representations in the lateral frontal pole

**DOI:** 10.1101/2021.07.28.453895

**Authors:** RC Lapate, IC Ballard, MK Heckner, M D’Esposito

**Affiliations:** University of California, Santa Barbara; University of California, Berkeley; Juelich Institute

## Abstract

Emotional states provide an ever-present source of contextual information that should inform behavioral goals. Despite the ubiquity of emotional signals in our environment, the neural mechanisms underlying their influence on goal-directed action remains unclear. Prior work suggests that the lateral frontal pole (FPl) is uniquely positioned to integrate affective information into cognitive control representations. We used pattern similarity analysis to examine the content of representations in FPl and interconnected mid-lateral prefrontal and amygdala circuitry. Healthy participants (n=37; n=21 females) were scanned while undergoing an event-related Affective Go/No-Go task, which requires goal-oriented action selection during emotional processing. We found that FPl contained conjunctive emotion-action goal representations that were related to successful cognitive control during emotional processing. These representations differed from conjunctive emotion-action goal representations found in the basolateral amygdala. While robust action goal representations were present in mid-lateral prefrontal cortex, they were not modulated by emotional valence. Finally, converging results from functional connectivity and multivoxel pattern analyses indicated that FPl’s emotional valence signals likely originated from interconnected subgenual ACC (BA25), which was in turn functionally coupled with the amygdala. Thus, our results identify a key pathway by which internal emotional states influence goal-directed behavior.

**Significance statement:** Optimal functioning in everyday life requires behavioral regulation that flexibly adapts to dynamically changing emotional states. However, precisely how emotional states influence goal-directed action remains unclear. Unveiling the neural architecture that supports emotion-goal integration is critical for our understanding of disorders such as psychopathy, which is characterized by deficits in incorporating emotional cues into goals, as well as mood and anxiety disorders, which are characterized by impaired goal-based emotion regulation. Our study identifies a key circuit through which emotional states influence goal-directed behavior. This circuitry comprised the lateral frontal pole (FPl), which represented integrated emotion-goal information, as well as interconnected amygdala and subgenual ACC, which conveyed emotional signals to FPl.

## Introduction

Optimal functioning in everyday life requires behavioral control that is goal oriented yet flexible to dynamically changing contexts. One’s emotional state—and the emotional states of others—are a primary source of such context. Emotional cues, such as others’ facial expressions, often provoke automatic action tendencies, such as approach towards appetitive stimuli, and avoidance of aversive stimuli. These emotion-driven action tendencies can inform, amplify, or interfere with goal-directed behavior. For instance, receiving a welcoming smile in a new circle of neuroscientists potentiates the existing (approach-related) goal pursuit of sharing a collaborative research proposal (whereas a scorn would likely provoke the opposite reaction, hindering goal completion). Despite the ubiquity of emotional information in our environment, the neural mechanisms underlying the potentiation (or interference) of goal-directed behavior by emotional signals remain underspecified.

Recent anatomical evidence suggests that the lateral frontopolar cortex (FPl) may mediate the influence of emotion on goal-directed behavior. FPl is ideally positioned to integrate and transmit information about internal states to the rest of lateral prefrontal cortex (LPFC). Unlike other LPFC regions, FPl receives projections from the amygdala via the ventral amygdalofugal pathway (Kamali et al. 2016; Folloni et al. 2019). Stronger structural connectivity of this pathway is associated with a stronger influence of emotion on action execution (Bramson et al. 2020). Disruption of FPl with transcranial magnetic stimulation amplifies the influence of emotional expressions on approach and avoidance behavior (Volman et al. 2011). Microstructural properties of the frontopolar cortex—including relatively low cell body density and reduced laminar differentiation, combined with increased dendritic length and spine number, compared to caudal and mid-lateral PFC regions—suggest that it has the highest level of information integration within the PFC (Jacobs et al. 2001; Badre and D’Esposito 2009; Badre and Nee 2018). However, because prior work has not employed multivariate methods that measure FPl representations, it remains unclear whether FPl’s goal representations differ depending on ongoing emotional states (i.e., reflecting emotion-goal integration). Neural representations that integrate across distinct dimensions, such as goals and emotional states (also called conjunctive representations) have been postulated to be particularly advantageous for flexible and context-sensitive behavior, which is critical during emotional processing (Badre et al. 2021; Fusi, Miller, and Rigotti 2016).

In order to test whether FPl representations integrate across goals and emotional information and are associated with the influence of emotional signals on goal-directed behavior, we adapted a task that requires cognitive control during emotional processing, the Affective Go/No-Go (AGNG) task, for an event-related fMRI experiment. In the AGNG task, emotional valence (“Positive” vs. “Negative”, here happy and fearful faces) and goal (i.e., action goals: “Go” vs. “No-Go”) are manipulated orthogonally (**Fig. 1A**). Critically, emotion biases action goals depending on emotion-action congruency: In some trials, positive stimuli are “Go” targets (emotion-action congruent); in others, negative stimuli are “Go” targets (emotion-action incongruent), and vice-versa. Positive stimuli typically facilitate approach behavior (e.g., reducing “Go” reaction times (RTs)), whereas negative stimuli facilitate avoidance (i.e., increasing “No-Go” accuracy). Thus, “Go” RT and “No-Go” accuracy difference scores in the AGNG task provided indices of emotion-driven influence on action, i.e., affect-to-motor spillover.

**Figure 1.**
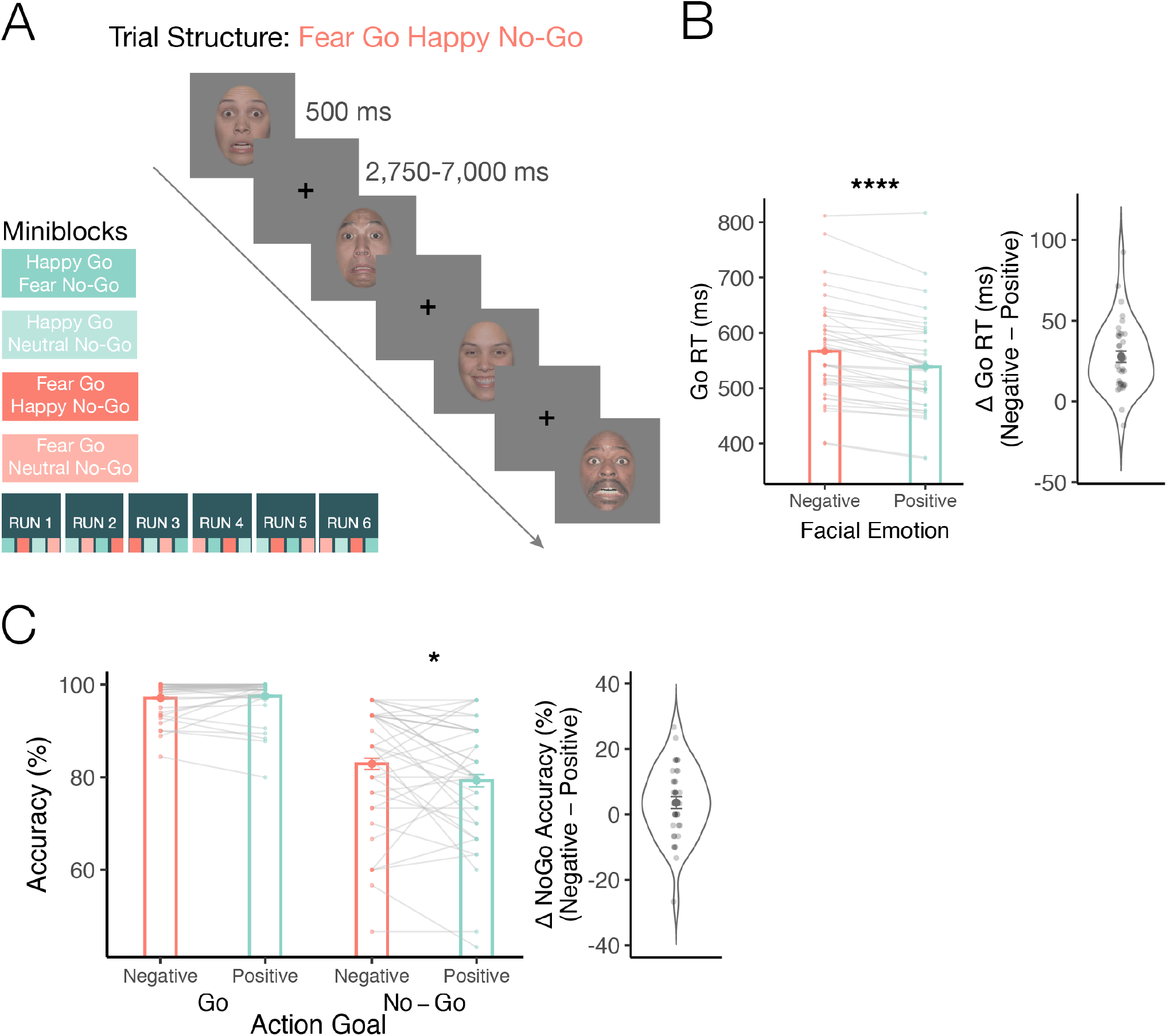
Emotional valence biases action. **(A)** The trial structure of the Affective Go/No-Go Task is shown. At the start of each condition miniblock (n=20 trials), participants were asked to press a button (“Go”) in response to either happy or fearful faces, and to withhold responses (“No-Go”) following the presentation of non-target emotional expressions (happy, fearful or neutral faces). The experiment comprised a total of six fMRI scans, and each fMRI scanner run contained n=4 miniblocks presented in counterbalanced order. **(B)** Reaction time (RT) data for “Go” trials. Positive emotional valence (happy facial expressions) facilitates approach responses compared to negative valence (fearful expressions), reducing “Go” reaction times. The violin plot shows the distribution of the valence difference score in RTs (Negative – Positive). **(C)** Task accuracy data. Negative emotional valence facilitates avoidance responses, increasing accuracy for “No-Go” trials. The violin plot shows the distribution of the valence difference score in No-Go accuracy (Negative – Positive). *\*p <* .*05 **** p<* .*0001*

We interrogated the content of neural representations in FPl and interconnected brain regions, including mid-LPFC (known to represent task goals (Waskom et al. 2014; Cole, Ito, and Braver 2016)) and basolateral amygdala (known to support emotional valence encoding (Tye 2018)). In order to test whether emotion modulated action goal representations, we employed pattern similarity analysis (Kriegeskorte and Kievit 2013). Classifier decoding and functional connectivity analyses complemented pattern similarity analysis to further reveal the informational structure and behavioral correlates of emotion and action goal representations in FPl and interconnected circuitry. We hypothesized that FPl would represent integrated emotion and action goals; conversely, that basolateral amygdala and mid-LPFC would contain separate emotion and goal representations, respectively.

## Materials and Methods

### Participants

Forty participants were recruited from Berkeley, CA (M = 22.22; SD = 3.14; range = 18-29; 24 female). Two subjects were excluded due to excessive motion (>3mm) and one chose not to complete the study. Thus, the full sample analyzed here comprised 37 participants (M = 22.24; SD = 3.22; range = 18-29; 21 female). All participants were healthy, with no self-reported history of neurological or psychiatric disorders, and had normal or corrected-to-normal visual acuity. Written informed consent was obtained from each subject. Subjects were recruited at the University of California, Berkeley. All study procedures were approved by the UC Berkeley Committee for the Protection of Human Subjects. Participants were compensated monetarily for their participation.

### Procedure

#### Overview

Participants underwent the Affective Go/No-Go task (AGNG) in the fMRI scanner as part of a longitudinal transcranial magnetic stimulation study on prefrontal mechanisms of emotion regulation. As part of this larger study, participants performed a resting state and diffusion imaging scan (data not reported here). Following the informed consent procedure, participants practiced a few trials of the AGNG task before beginning the experiment. After approximately 40 minutes of fMRI data collection during the AGNG task, a high resolution T1-weighted anatomical scan was obtained. At the end of the experiment, participants completed questionnaires (data not reported here).

#### Affective Go/No-Go (AGNG) task

In the MRI scanner, participants completed the AGNG task. The task was comprised of six functional runs (lasting approximately 7min/each). Each functional run contained 4 action goal + emotion target miniblocks: “*Go Happy, No-Go Fear*”, “*Go Fear, No-Go Happy*”, “*Go Happy, No-Go Neutral*” and “*Go Fear, No-Go Neutral*”. Each miniblock contained 20 trials, 75% (15/20) of which were “Go” trials, and 25% (5/20) were “No-Go” trials. Before each miniblock, participants were instructed to press a button on a handheld button box for faces that matched the “Go” condition of the miniblock, and to withhold pressing the button for faces that matched the “No-Go” condition of the miniblock. Those four miniblocks were presented in counterbalanced orders across the 6 functional runs (see **Fig. 1A**), and 2 scan run orders were used (counterbalanced across subjects).

In each trial, a fixation cross appeared for 500ms, followed by a face image that was either a target (“Go”) or nontarget (“No-Go”), which was presented for 500ms. Then, a 2750-7000ms inter-trial interval ITI followed (sampled from an exponential distribution). The AGNG task totaled 80 trials/run (n=480 trials total across the task) and took approximately 40 minutes to complete.

#### Face Stimuli

Emotional faces (happy, neutral and fearful) consisted of 12 identities (half female) selected from the Macbrain Face Stimulus Set http://www.macbrain.org/resources.htm). Faces were cropped to remove hair and neck, matched for average luminance and RMS contrast. Emotional faces were presented at 13º x 13º using PsychoPy (Peirce et al. 2007). The full list of emotional face stimuli, example stimuli, and stimulus presentation scripts used in this study are available online at https://osf.io/rqa8w/.

#### AGNG metrics & behavioral analyses

As dependent measures, we examined task accuracy, as well as two indices of the emotion-driven influence on task behavior—i.e., “affect-to-motor spillover”. Emotional valence typically biases behavioral action in a valence congruent manner, whereby appetitive stimuli (such as happy faces) facilitate approach behavior (“Go” responses, as reflected by shortened reaction times), whereas aversive stimuli (such as fearful faces) typically facilitate avoidance (increasing accuracy in “No-Go” trials). Therefore, we computed 2 difference scores reflecting the magnitude of affect-to-motor spillover; in “Go” trials, the reaction time difference between “Go Fear” and “Go Happy” trials (Fear – Happy RT); and for “No-Go” trials, the accuracy difference score between “No-Go Fear” and “No-Go Happy” trials (No-Go Fear – Happy Accuracy). We examined whether RT and accuracy were modulated by emotional valence using the lme4 package (Bates et al. 2015), anova and emmeans (https://github.com/rvlenth/emmeans) functions in R.

### Functional MRI Methods

#### Image Acquisition

Neuroimaging data were acquired in the UC Berkeley Henry H. Wheeler, Jr. Brain Imaging Center with a Siemens TIM/Trio 3T MRI scanner with a 32-channel RF head coil. Whole-brain Blood Oxygen Level-Dependent (BOLD) functional Magnetic Resonance Imaging (fMRI) data were obtained using a T2*-weighted 2x accelerated multiband echo-planar imaging (EPI) sequence (52 axial slices, 2.5 mm^3^ isotropic voxels; 84 × 84 matrix, TR= 2000 ms; TE= 30.2 ms; flip angle = 80°; 222 image volumes per run). High-resolution T1-weighted MPRAGE gradient-echo sequence images were collected at the end of the session for spatial normalization (176 × 256 × 256 matrix of 1 mm^3^ isotropic voxels; TR = 2300 ms; TE = 2.98 ms; flip angle = 9°).

#### fMRI data preprocessing

Functional neuroimaging data were preprocessed using FEAT (Smith et al. 2004; Jenkinson et al. 2012) implemented in FSL version 6.0.1. Preprocessing steps included removal of the first four functional volumes, high-pass filtering with a 90 s cutoff, FILM correction for autocorrelation in the BOLD signal, slice-time correction, motion correction using MCFLIRT, and creation of a confound matrix of points of framewise displacement changes greater than 0.5mm to be used as regressors of non-interest in the analyses to control for movement-confounded activation. Data were smoothed with using a 3 mm full width at half maximum (FWHM) Gaussian spatial filter. Functional images were co-registered to a high-resolution (T1-weighted) anatomical image using a linear rigid body (6-DOF) transform (while maintaining native functional resolution, i.e., 2.5 mm^3^ isotropic).

#### Regions of Interest (ROIs)

##### Subcortical

The basolateral amygdala ROI was defined using the Tyszka et al. CITI atlas basolateral nuclear group definition (i.e., lateral, basolateral and basomedial/accessory basal nuclei) thresholded at 50% (Tyszka and Pauli 2016) and registered from MNI to participants’ structural space using FNIRT (10 mm warp resolution) while maintaining native resolution (2.5 mm^3^ isotropic).

##### Cortical

Prefrontal ROIs (FPl, mid-LPFC (BA9-46), BA25/subgenual ACC and BA32) were obtained from the Oxford PFC consensus atlas (Neubert et al. 2014; Sallet et al. 2013; Verhagen 2018), thresholded at 25%, and registered to participants’ native surface space using Freesurfer (Reuter et al. 2012). For the mid-LPFC ROI, we focused on 9-46 (9-46d + 9-46v) because this mid-LPFC region, located in the middle frontal gyrus, has been shown to be a critical site of convergence of information relevant for cognitive-control (Derek Evan Nee and D’Esposito 2016). For the ventromedial PFC ROIs, we focused on BA25 & BA32 because both of these regions are known to be interconnected with the basolateral amygdala and frontopolar cortex in the non-human primate (Barbas and Pandya 1989; Joyce and Barbas 2018; Medalla and Barbas 2010). Vertex coordinates in each of these ROIs were transformed into the native anatomical (volumetric) space, and ROI masks in volumetric space were constructed by projecting half the distance of the cortical thickness at each vertex, requiring that a functional voxel be filled at least 50% by the label, and labeling the intersected voxels.

#### fMRI data modeling

For multivariate analyses (pattern similarity & classifier analyses), we obtained voxel-and trial-wise BOLD activation parameters estimates using the Least-Squares All (LS-A) GLM approach (Mumford et al. 2012) and FEAT modeling in FSL (Jenkinson et al. 2012). Single trials were modeled using a canonical hemodynamic response function (Double γ). Only correct trials were included in the analysis reported here (error trials were included in the model as a nuisance regressor). This amounted to up to 480 trials per participant (Mean error = 5.783%, SD = 4.143%). In addition to single trial and error trial regressors, we modeled the 8s instruction epoch using a canonical hemodynamic response function. Parameter estimates extracted from each ROI were regularized with multivariate noise normalization (Walther et al. 2016). To do so, we obtained an estimate of the noise covariance from the residuals of the general linear models from each ROI. This matrix was then regularized using the optimal shrinkage parameter, inverted, and multiplied by the vector of betas for each trial (Ballard, Wagner, and McClure 2019; Walther et al. 2016). This approach removes nuisance correlations between voxels that arise due to physiological and instrument noise.

#### Pattern similarity analysis

We used pattern similarity analysis to examine the structure of emotion and action goal representations in FPl and interconnected circuitry (Kriegeskorte and Kievit 2013). Pattern similarity analysis tests whether the inter-trial similarity structure of multivoxel activity patterns (i.e., the *neural similarity matrix*) is explained by experimental factors – in the current experiment, emotional valence, action goal, and their interaction – which are expressed as pattern similarity analysis template matrices (**Fig. 2**). To obtain the neural similarity matrix for each participant and ROI, we computed pairwise Pearson correlations between pairs of multivoxel patterns obtained from all trials *except for* trials from the same run (i.e., a between-runs correlation approach, which is akin to a leave-one-run-out cross validation approach for multivariate classification analyses, and minimizes inflated correlations due to data dependencies and temporal autocorrelation (**Fig 2A**)). Pearson’s correlations are a distance metric that are invariant to scale changes in multivoxel patterns. This approach yielded a trial-wise neural similarity matrix for each participant and ROI.

**Figure 2.**
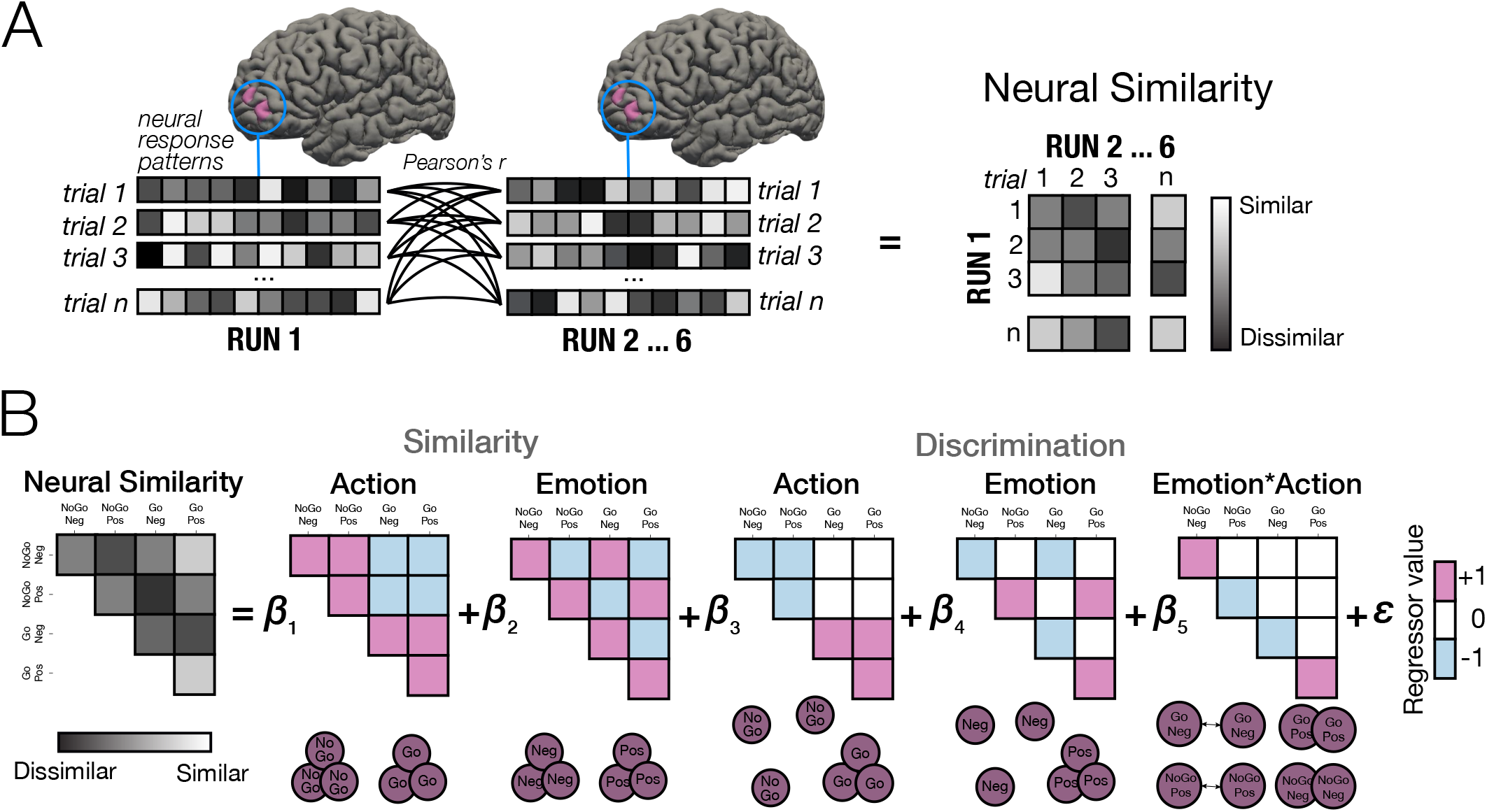
Pattern similarity analysis strategy. **(A)** Neural representational similarity matrices were obtained for each ROI by computing the correlation of multivoxel activity patterns across all trials of each condition obtained from independent scans (i.e., between-runs pattern similarity analysis). Each box denotes a voxel in the ROI, and each row denotes a trial. **(B)** Next, we used condition-specific template matrices to test whether neural similarity matrices were explained by action goal, emotional valence, and/or their interaction (conjunctive emotion*action goal representation). Template matrices tested differential representational distances by condition (*discrimination* matrices) as well as equivalent representational distances across different conditions (*similarity* matrices). Lower circles depict the representational structure captured by each regressor.

Next, we fit a multiple regression model using condition-specific template matrices to test whether the similarity structure of multivoxel patterns in each ROI’s neural similarity matrix was modulated by emotional valence, action goal, and/or their interaction (**Fig 2B**). Because the AGNG task captures the *differential* impact of emotion on action based on action-goal congruency (i.e., positive and negative emotion differentially modulate “Go” and “No-Go” responses) our primary set of model matrices tested for *discriminative* similarity structures across unique combinations of emotion and action goal levels. In other words, these discriminative model matrices assess whether representational distances across a high-dimensional space are *differentially* modulated by emotion type or action-goal type (for instance, whether the average representational distance amongst distinct positively valenced events (trials) differs from the distance amongst negatively valenced events, or whether the distance amongst “No-Go” patterns differed from that of “Go” patterns). Critically, the *interactive* discriminative matrix of emotional valence and action goal regressors (hereafter referred to as the emotion*action regressor) captures the emotion-action congruency phenomenon studied here: “*Go Positive*” and “*No-Go Negative*” representations (i.e., congruent emotion–action conditions) are modeled closer in a high-dimensional space compared to “*No-Go Positive*” and “*Go Negative*” representations (incongruent conditions). Thus, by using three orthogonal discrimination matrices, we simultaneously modeled the impact of emotional valence, rule, and their interaction (conjunction) on neural similarity matrices (**Fig. 2B**).

To control for similarities across trials attributable to shared emotional valence or action goal conditions, we also included *similarity* template matrices in our simultaneous regression model (**Fig. 2B**). Those *similarity* matrices, in contrast to the *discrimination* matrices, tested for within-condition inter-trial similarity while assuming *equivalent* representational distances across conditions (e.g., equivalent average pattern distance for positively and negatively valenced trials, collapsing across Go/NoGo action goal conditions). We fit the neural similarity matrices to these five total template matrices using a multiple regression mixed-model framework, where subject was modeled as a random factor, and each unique template matrix was included in the subject error term.

This allowed us to test whether FPl and interconnected amygdala and LPFC circuitry represented the interaction of emotional valence or action goal (i.e., conjunctive representations), vs. whether it represented emotional valence and/or action goal dimensions (*Positive vs. Negative*; *Go vs. No-Go*) independently of one other. Importantly, this approach also allowed us to examine interactive effects while controlling for potential motor confounds arising from the “Go” vs “No-Go” conditions, given that the similarity patterns of “Go” and “No-Go” are entered simultaneously in the same regression model.

#### Multivariate pattern analysis (MVPA)

We used MVPA to examine whether emotional valence (*Positive vs. Negative*) and action goal (*Go vs. No-Go*) were linearly decodable in FPl and interconnected circuitry. MVPA used multivariate-noise normalized single-trial betas as described above and was implemented with Nilearn (Abraham et al. 2014). For each subject and ROI, we used a multivariate logistic regression model *(l2 penalty; C=1)* to iteratively train the classifier on z-scored data. We assessed classifier performance using a leave-one-run-out cross validation scheme. Classification performance was evaluated using the area under the curve (AUC) (i.e., where 0.5 is chance performance). We examined whether the logistic classifier could distinguish emotional valence classes (*Positive vs. Negative*) as well as action goal (*Go vs. No-Go*). We used Nilearn’s parameter (class_weight=‘balanced’) to automatically adjust weights according to class frequencies in the input data (which is important for classification of *Go vs. No-Go* classes). To examine whether classification accuracy differed from chance, we combined run-wise classifier AUCs (‐0.5) across subjects using a mixed-model approach, where subject and run were entered as random factors, and tested whether the intercept differed significantly from 0.

#### MVPA-behavior correlations

We tested the behavioral relevance of the strength of classifier evidence for emotional valence and action goal in FPl using both an intra-individual analysis and an across-subjects analysis. For the intra-individual analysis, we regressed run-wise estimates of classifier AUC for emotional valence and action goal on run-wise estimates of task accuracy and affect-to-motor spillover (i.e., emotion-based difference scores in RT (“Go” trials) and accuracy (“No-Go” trials)). For analyses across subjects, we examined the Spearman’s rho coefficient of the association between classifier AUC, task accuracy and affect-to-motor spillover indices. Difference of dependent correlation coefficients were tested using the cocor package in R (http://comparingcorrelations.org/)

#### Functional connectivity analysis

To examine the putative origins of emotional valence information arriving in FPl, we used psychophysiological interaction analysis (PPI, (O’Reilly et al. 2012)). To do so, we first extracted the mean time series from FPl and basolateral amygdala (BLA) regions of interest (ROIs). We then ran a separate FEAT analysis that included emotional valence task regressors, the demeaned timecourse of the ROI seed, as well as the interaction between this timecourse and the regressors encoding positive and negative emotional stimuli. The betas (parameter estimates) obtained from the interaction contrast were extracted for each subject, functional run, and ROI. As extreme outliers were observed, run-wise betas that exceeded +/-4 SD from the mean (across subjects) were excluded prior to analyses. To test whether BLA-BA25 significantly coupled during the AGNG task, betas obtained were entered into a mixed effects model (where subject and run were entered as random factors) to test whether the intercept differed significantly from zero. Finally, run-wise variation in BA25<->FPl functional coupling (PPI betas) were used to predict emotional valence classifier accuracy in FPl (with subject and run entered as random factors).

## Results

### Behavioral analysis: Emotion biases action goal

We first verified whether emotion biased the execution of action goals in the AGNG task. As predicted, we found that emotion-action congruency influenced behavior: In “Go” trials, “Go” target happy faces reduced reaction times relative to “Go” fearful faces (*F*(1,36) = 63.047, *p* < 0.001; **Fig. 1B**), which is consistent with appetitive states facilitating one’s behavioral approach in the goal-congruent “Go” condition. In contrast, fearful faces increased accuracy in “No-Go” trials compared to happy faces (*Z* = 2.024; *p* = 0.043; **Fig. 1C**), which is consistent with aversive states facilitating avoidance in the goal congruent “No-Go” condition. The impact of emotion on AGNG accuracy was specific to “No-Go” trials, as indicated by the interaction of emotional valence and action goal (*F*(1,36) = 4.104, *p* = 0.05). In sum, emotional valence facilitated *or* hindered action depending on the congruency of emotion and action goal, with positive emotional states facilitating approach (“Go”) and negative emotional states facilitating avoidance (“No-Go”) responses.

### Pattern Similarity Analysis: Conjunctive representations of emotion and action goal in FPl and amygdala

Next, we tested if emotion-action congruency modulated the structure of neural representations in FPl and interconnected circuitry. In other words, do emotional states change the representation of action goals? If so, this implies that a region carries a conjunctive (i.e., integrated) representation of emotion and goal. To answer this question, we used pattern similarity analysis. Pattern similarity analysis tests whether the inter-trial similarity structure of multivoxel activity patterns (i.e., the neural similarity matrix) is explained by experimental factors – in this case, emotional valence, action goal, and, critically, their interaction (**Fig. 2**).

To that end, we first computed neural similarity matrices for each ROI (**Fig. 2A**). Next, we used condition-specific template matrices to test whether neural similarity matrices were explained by emotional valence, action goal, and/or their interaction (i.e., conjunctive emotion*action representations). To account for the distinct ways in which emotion and action may be represented in the brain, we used two sets of template matrices in our model: (1) One that tested for differential representational distances between different emotion and action goal conditions, as well as their interaction (emotion*action). These comprised our primary regressors of interest (we refer to these matrices as condition *discrimination* matrices). (2) We also included orthogonal template matrices that tested for within-condition similarity (equivalency) in representational distances across emotion and action goal conditions (*similarity* matrices) (**Fig. 2B**). (see *Methods* for additional details). We fit these template matrices on the neural similarity matrix using a simultaneous multiple regression mixed-model approach, with subject as a random intercept, and condition (each template matrix model) included in the subject error term. This approach tests whether FPl and interconnected circuitry have integrated or independent emotion/action goal representations.

We found evidence of conjunctive representations of emotion and action goal in FPl, as demonstrated by the significant fit of the interaction emotion*action regressor (*B* = 0.003 (*SE* = 0.001), *t* = 2.354, *p* = 0.024; **Fig. 3A**). This result indicates that trials were represented more similarly in a high-multidimensional space when emotion and action goal were congruent (“*Go Positive*” and “*No-Go Negative*”) than incongruent. Thus, emotional context sculpted action goal representations in FPl, suggesting that FPl integrates internal states and goal representations. In contrast, in mid-LPFC, the representation of action goals was unaltered by emotional valence (emotion*action interaction regressor *p* > 0.64; **Fig. 3B**). Only action goal significantly explained the neural similarity structure observed in mid-LPFC (action goal similarity *B* = 0.001 (*SE* = 0.0004), *t* = 3.194, p < 0.003; action goal discrimination *B* = -0.005 (*SE* = 0.001), *t* = -3.364, *p* < 0.002). Conjunctive representations of emotional valence and action goal were significantly stronger in FPl than in mid-LPFC (*Z* = 5.071 *p* < 0.0001), suggesting regional specificity in conjunctive emotion-action representations in LPFC circuitry. We note that the emotional valence discrimination regressor also explained significant variance in FPl’s similarity structure (*B* = -0.002 (*SE* = 0.001), *t* = -2.124, *p* = 0.041). However, caution is warranted when interpreting simple effects from lower order terms in a model that includes an interaction (accordingly, the simple effect of valence in FPl is non-significant when the interaction of emotion*action is excluded from the model (*p* > 0.79), suggesting that the representational geometry of emotional valence in FPl may depend on goal state).

**Figure 3.**
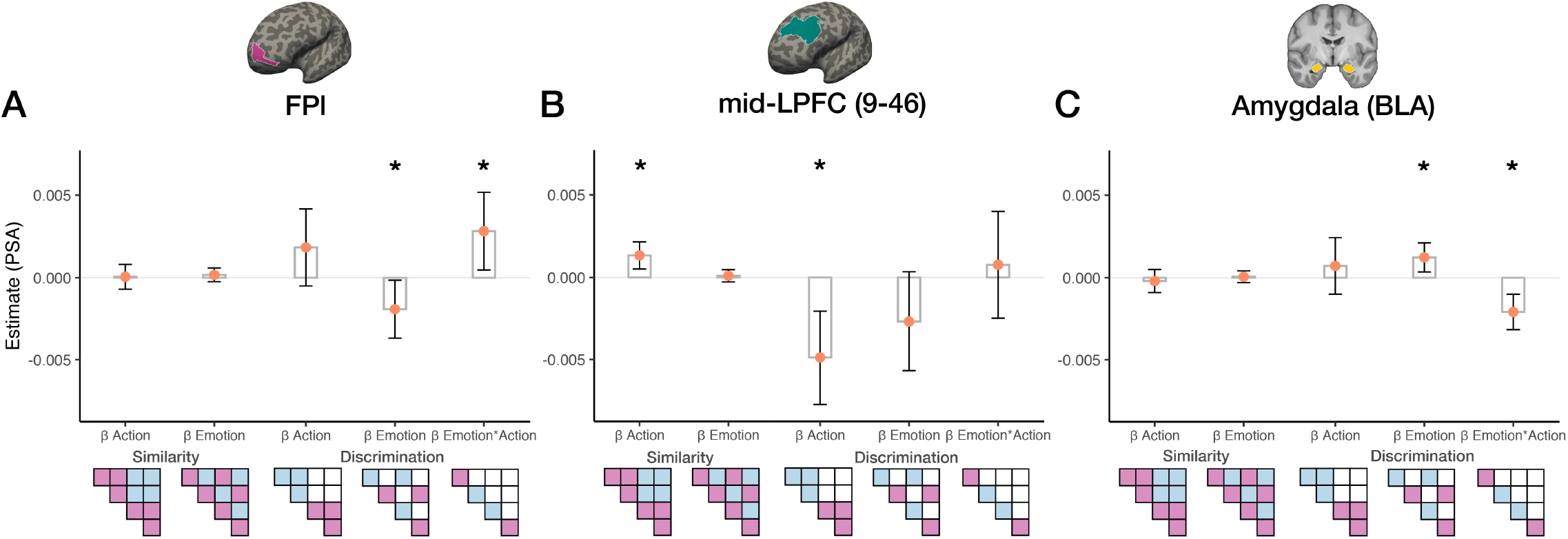
Pattern similarity analysis (PSA). A simultaneous regression analysis of orthogonal template matrices on neural similarity matrices using a mixed-effects model revealed that **(A)** FPl and **(C)** basolateral amygdala (BLA) show evidence of conjunctive emotion*action goal representations whereas **(B)** mid-LPFC shows evidence of action goal representations unaltered by emotional valence. The y-axis shows the fixed-effects regression weights from the mixed-model regression for each ROI. While emotion and action goal are integrated into a higher-order representation in both FPl and BLA, their representational geometries are flipped in sign. Specifically, emotion-action congruent conditions (such as *Go Positive*, or *No-Go Negative*) are represented closer in a high-dimensional space in FPl relative to incongruent conditions (such as *Go Negative*, or *No-Go Positive*) **(A)**, whereas the opposite was observed in the amygdala, wherein emotion-action incongruent trials were represented more similarly than congruent trials **(C)**. The strength of emotion-action conjunctive coding differed significantly across the three ROIs, being significantly stronger in FPl and amygdala compared to mid-LPFC, *p*s < 0.0001. *\*p < 0*.*05*

Next, we examined the representational structure of emotion and action goal in the basolateral amygdala. We found that the basolateral amygdala, like the FPl, also expressed conjunctive emotion-action goal representations (*B* = -0.002 (*SE* = 0.001), *t* = -3.797, *p* < 0.001; **Fig. 3C**). However, the conjunctive emotion*action goal coding in the amygdala differed markedly from that in FPl: in the amygdala, emotion-action *incongruent* trials were represented closer together in a high-multidimensional space relative to emotion-action congruent trials. Conjunctive emotion*action goal representations were stronger in the amygdala than in mid-LPFC (*Z* = -10.400, *p* < 0.0001). As expected, given their opposite signs, conjunctive coding in FPl significantly differed from that observed in the amygdala (*Z* = 6.312, *p* < 0.0001). We note that the emotional valence discrimination regressor also explained significant variance in amygdalar similarity structure (*B* = 0.001 (*SE* = 0.0004,) *t* = 2.742, *p* < 0.009), which, as mentioned above, should be interpreted with caution given that the emotion*action interaction is significant (as was the case with FPl, the simple effect of emotional valence in the amygdala was not observed when the emotion*action regressor was removed from the model, *p* > 0.46). In summary, emotional valence shaped the representation of action goal in the FPl-amygdala circuitry, with regionally specific representational geometries.

We next tested whether the strength of conjunctive emotion*action goal representations correlated with task performance. We hypothesized that the conjunctive representational structure observed in FPl, in which emotion-congruent trials are represented more similarly, is behaviorally advantageous. We found that conjunctive coding in FPl was associated with higher task accuracy (*Spearman’s rho* = 0.33, *p* = 0.048; **Fig. 4**). This relationship was driven by the similarity amongst congruent emotion-action trials (*rho* = 0.42, *p* = 0.01) (as opposed to dissimilarity amongst incongruent trials, *rho* = -0.1, *p* > 0.5). This association was not observed in the basolateral amygdala (*rho* = 0.12, *p* > 0.47); although note that FPl vs. amygdala correlations were not significantly different (*p* > 0.1). These results suggest that conjunctive emotion-action representations in FPl support adaptive action selection during emotional processing.

**Figure 4.**
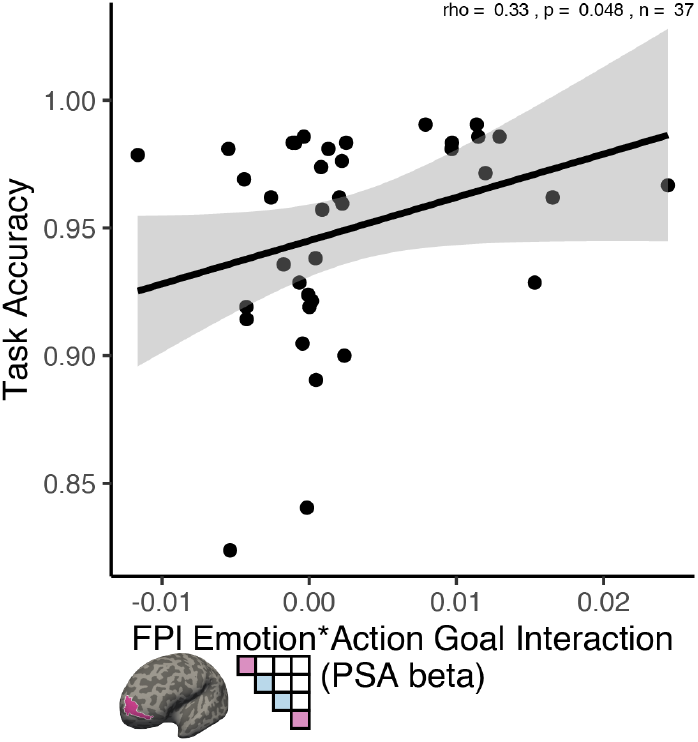
Association between task accuracy and the magnitude of conjunctive representation of emotion and action goal in FPl (as indicated by the beta coefficient of the emotion*action goal interaction in the pattern similarity analysis (PSA) regression model shown in 3A) across participants.

### The behavioral relevance of action goal and emotional valence representations in FPl

High dimensional (conjunctive) cognitive control representations are useful because they facilitate context-appropriate behavior, but perfectly conjunctive representations carry an information cost and a resulting lack of flexibility to adapt to changing task demands (Fusi, Miller, and Rigotti 2016; Badre et al. 2021; Barak, Rigotti, and Fusi 2013). For example, if representations of emotion and action goal are perfectly conjunctive, then a downstream neural population will be unable to distinguish emotional valence irrespective of action goal. In other words, there is a benefit to control representations that are both conjunctive and linearly separable (Bernardi et al. 2020). Thus, we examined whether emotional valence and action goal in FPl and interconnected circuitry were linearly separable. We used and a leave-one-run out cross validation approach, and combined run-level logistic classifier AUCs across subjects using mixed-models, to evaluate the discriminability of representations of emotional valence and action goal.

In FPl, we found evidence of linearly separable representations of *both* emotional valence and action goal (emotional valence classifier accuracy *M* = 51.6%, *B* = 0.016 (*SE* = 0.007), *t* = 2.223, *p* = 0.032; action goal classifier accuracy *M* = 51.99%, *B* = 0.0198 (*SE* = 0.007), *t* = 2.84, *p* = 0.015, respectively; **Fig. 5A**). In mid-LPFC, classifier evidence was above chance for action goal (*M* = 57.11%, *B* = 0.07 (*SE* = 0.012), *t* = 5.737, *p* < 0.001) but not emotional valence (*M* = 50.48%, *B* = 0.005 (*SE* = 0.007), *t* = 0.642, *p* > 0.53) (**Fig. 5B**). In the basolateral amygdala, neither emotional valence nor action goal categories were discriminable (valence *M* = 50.66%, *p* > 0.39, and *M* = 50.01%, *p* > 0.99, respectively; **Fig. 5C**). The strength of classifier evidence for action goal differed across regions (*F* = 49.361, *p* < 0.001), such that action goal representations in both mid-LPFC and in FPl were significantly stronger than in the basolateral amygdala (*t* = -9.613, *p* < 0.0001 and *t* = -2.679, *p* = 0.008, respectively), and greater in mid-LPFC than in FPl (*t* = 6.934, *p* < 0.0001). However, the strength of classifier evidence for emotional valence representation, while only significant in FPl, did not differ significantly across regions (*p*s > 0.134).

**Figure 5.**
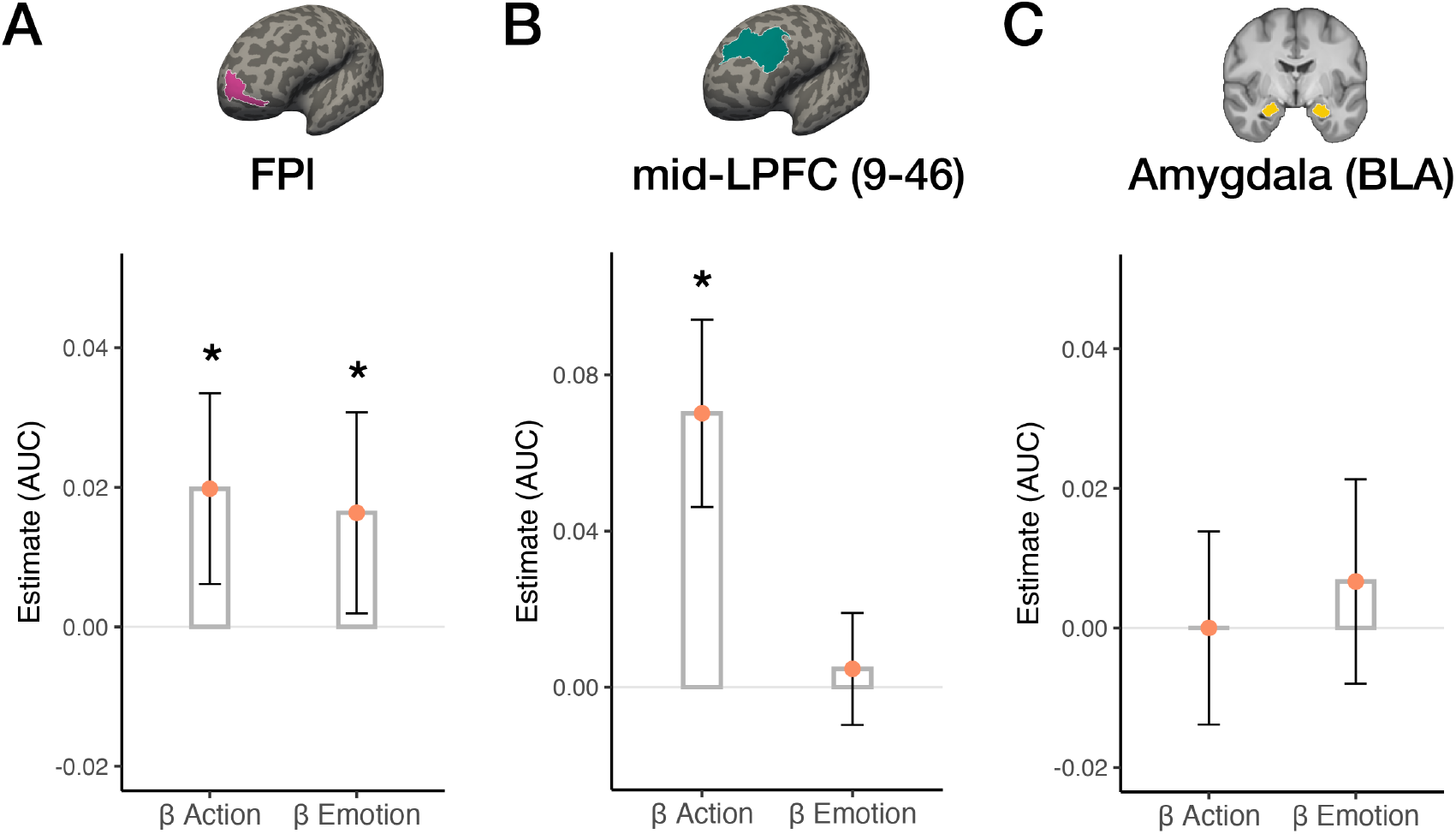
Decoding of emotional valence and action goal representations in FPl-amygdala circuitry. Subject- and run-wise classifier AUC for emotional valence and action goal were tested against zero using a mixed-effects model for each ROI; beta coefficients for each regression model are plotted for **(A)** FPl, **(B)** mid-LPFC and **(C)** Amygdala (BLA). *\*p <* .*05*

Given that emotional valence and action goal representations in FPl were linearly separable, we next examined whether they were associated with task performance and emotion-driven behavior in the AGNG task. Stronger action goal representations in FPl’s were associated with greater task accuracy within individuals and across runs (mixed model *B* = 0.077 (*SE* = 0.033), *t* = 2.322, *p* = 0.021; **Fig. 6A**). Across individuals, and collapsing across runs, this association was also positive, albeit non-significant (*rho* = 0.29, *p* = 0.08; **Fig. 6B**). Moreover, stronger action goal representations in FPl were associated with less affect-to-motor spillover across participants, as indexed by slower reaction times in negatively valenced “Go” trials relative to positively valenced “Go” trials (*rho* = -0.38, *p* = 0.02; **Fig. 6C**). Conversely, stronger emotional valence representations in FPl correlated with *increased* affect-to-motor spillover in “No-Go” trials, as indexed by higher accuracy in negatively valenced “No-Go” relative to positively valenced “No-Go” trials (*rho* = 0.37, *p* = 0.02; **Fig. 6D**). In other words, emotion-driven action tendencies benefited goal-driven action in trials where emotional-valence and goal-based action were congruent, as is the case whenever negative facial expressions were the “No-Go” cues. All other relationships between decoding and behavior were non-significant. These results suggest that high fidelity action goal representations in FPl facilitate task performance and reduce the influence of emotion on behavior, whereas stronger emotion representations in FPl increase the influence of emotion on behavior. However, we note the caveat that action and emotion decoding were each only related to one of our two affect-to-motor spillover measures (“Go” RT and “No-Go” accuracy, respectively).

**Figure 6.**
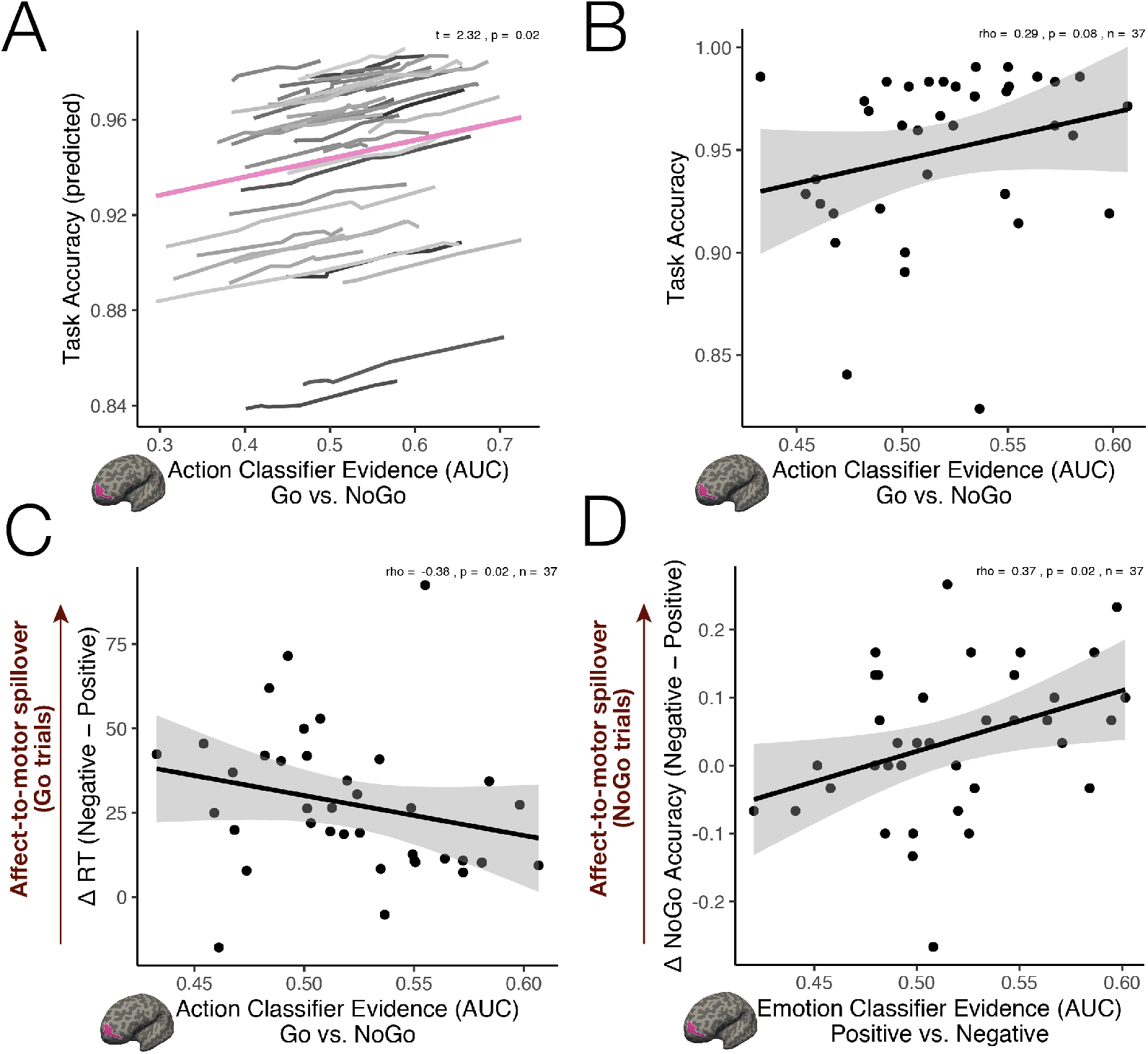
Association between behavior in the Affect Go/No-Go (AGNG) task and strength of action goal and emotional valence representations in FPl. **(A)** Stronger classifier evidence for action goal in FPl was associated with greater task performance within individuals, as evidenced by a mixed-effect model. **(B)** A similar trend was observed across subjects, wherein action-goal classifier evidence in FPl tended to correlate positively with task performance in the AGNG task. **(C)** Stronger classifier evidence for action goal in FPl was inversely associated with affect-to-motor spillover in Go trials, as indicated by the faster reaction times in Go-positive relative to Go-negative trials. **(D)** Conversely, stronger classifier evidence for emotion in FPl correlated with greater affect-to-motor spillover in No-Go trials, as indicated by higher accuracy in No-Go negative relative to No-Go positive trials.

### Medial prefrontal contributions to emotional valence representation in FPl

While emotional valence information was present in FPl and modulated action goal representations, it remains unclear how FPl gains access to this information. FPl’s access to internal states has been postulated to rely on projections from ventromedial PFC (vmPFC) regions (Badre and Nee 2018). Specifically, two distinct vmPFC regions, BA25 and BA32, are interconnected with the basolateral amygdala and project to the frontopolar cortex in the non-human primate (Barbas and Pandya 1989; Joyce and Barbas 2018; Medalla and Barbas 2010).

In order to probe whether vmPFC conveys emotional valence information to FPl, we tested whether the strength of emotional valence classifier evidence in FPl covaried between FPl and BA25 as well as between FPl and BA32. We found that the strength of emotional valence decoding in BA25 predicted emotional valence decoding in FPl (*B* = 0.152 (*SE* = 0.068), *t* = 2.219, *p* = 0.0275; **Fig. 7A**) (in contrast, emotional valence decoding in BA32 and FPl did not correlate, *p* > 0.27). Of note, BA25 is thought to be the primary vmPFC target and source of amygdala projections (Ghashghaei, Hilgetag, and Barbas 2007). Task-dependent functional connectivity analysis (psychophysiological interaction; PPI) confirmed that BA25 functionally coupled with the basolateral amygdala during negative emotional processing trials (*B* = 0.361 (*SE* = 0.139), *t* = 2.6, *p* < 0.001; **Fig. 7B**). (The PPI fit was nonsignificant in BA32 during negative or positive emotional processing (*p*s > 0.69), or in BA25 during positive emotional processing (*p* > 0.503)). Emotional valence classifier accuracy was above chance in BA25 (*M* = 51.95%, *B* = 0.02 (*SE* = 0.009), *t* = 2.195, *p* = 0.048), but not in BA32 (*M* = 49.27%, *B* = -0.007(*SE* = 0.008), *t* = -0.974, *p* = 0.336).

**Figure 7.**
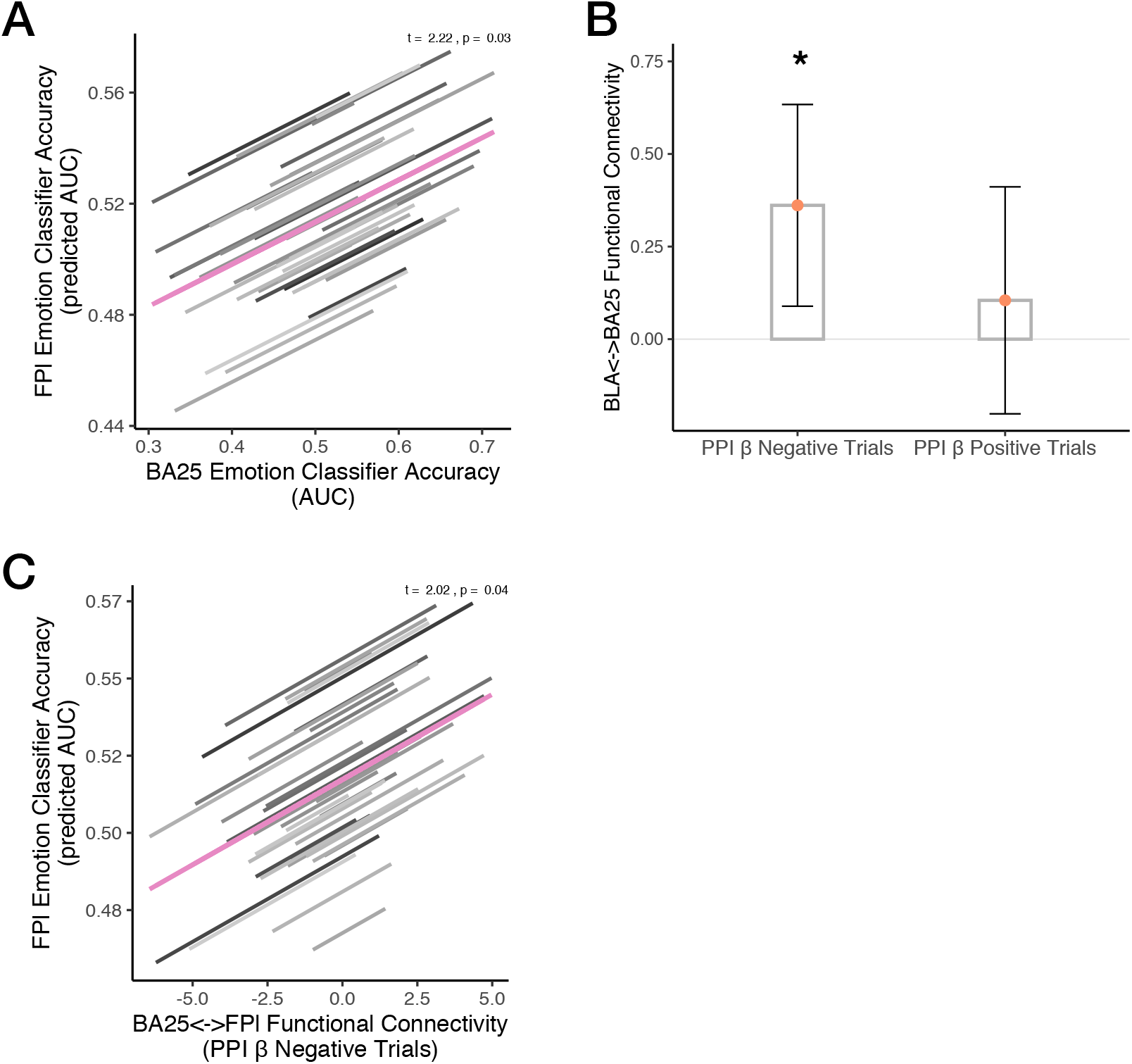
A pathway for emotional information flow from vmPFC (BA25) to FPl. **(A)** The strength of classifier evidence for emotional valence in BA25 (AUC) was a significant predictor of the strength of classifier evidence for emotional valence in FPl (AUC) (run-wise mixed model). **(B)** A psychophysiological interaction analysis (PPI) revealed that the basolateral amygdala (BLA) and BA25 functionally coupled during negative emotional processing trials, confirming well known dense anatomical projections between BLA and BA25 (Ghashghaei, Hilgetag, and Barbas 2007). **(C)** Further suggesting that BA25 may provide a key source of emotional valence information to FPl, the strength of BA25-FPl coupling during negative emotional processing (PPI) predicted the strength of classifier evidence for emotional valence in FPl (AUC).

We next examined whether functional interactions between BA25 and FPl were associated with FPl’s encoding of emotional valence. We found that the magnitude of BA25-FPl coupling during negative emotional processing correlated with the strength of emotional valence classifier evidence in FPl (*B* = 0.006 (*SE* = 0.003), *t* = 2.02, *p* = 0.045; **Fig. 7C**). Collectively, these data suggest that BA25 function facilitates the flow of emotional valence information into FPl, which contextualizes goal representations based on variation on ongoing emotional states.

## Discussion

Using pattern similarity analysis of fMRI data, we found that emotional context sculpted action goal representations in FPl: emotion and action goal were integrated in FPl into conjunctive representations. In contrast, in mid-LPFC, action goal representations were not modulated by emotion. In FPl, conjunctive emotion-action goal representations were modulated by emotion-action congruency. Moreover, the extent of these neural emotion-action congruency effects correlated with task accuracy. In FPl, but not in mid-LPFC or basolateral amygdala, action goal and emotional valence information were linearly separable, associated with task accuracy and with affect-to-motor spillover. Functional connectivity analyses revealed that BA25 (subgenual ACC) in the vmPFC served as the likely source of emotional valence information arriving in FPl. Collectively, these findings add to a growing literature pointing to a key role for FPl in the modulation of motivated action (Bramson et al. 2020; Volman et al. 2011; Kim et al. 2020). While prior work had demonstrated that FPl function plays an important role in the control of approach and avoidance behavioral tendencies (Bramson et al. 2020; Volman et al. 2011; Koch et al. 2018), the precise neural-representational mechanisms underlying the flexible utilization of goal-based signals and emotional context during cognitive control had not been previously specified.

### Implications for theories of cognitive control

Conjunctive representations have been postulated to both increase behavioral flexibility and minimize interference when behavior depends on context (Badre et al. 2021). These features should be particularly relevant in emotionally provocative situations: During emotional processing, contextual emotion-goal integration is critical—otherwise, behavioral responses may become excessively emotion driven (e.g., guided by automatic action tendencies such as approach or withdrawal irrespective of goals); or excessively goal driven, at the cost of failing to automatically incorporate important socio-emotional cues in the environment (as is the case in psychopathy; (Baskin-Sommers, Stuppy-Sullivan, and Buckholtz 2016)). Our finding that conjunctive emotion-action goal representations in FPl were associated with task performance supports the proposal that high-dimensional control representations are behaviorally advantageous (Fusi, Miller, and Rigotti 2016; Badre et al. 2021; Kikumoto and Mayr 2020; Bernardi et al. 2020).

The utility of conjunctive representations depends partially on the extent to which important components comprising the conjunction can be read out by downstream circuitry (Fusi, Miller, and Rigotti 2016; Badre et al. 2021). Accordingly, we found that conjunctive representations of emotional valence and action rule in FPl were linearly separable. Moreover, the strength of action goal representations in FPl correlated with task performance and reduced affect-to-motor spillover, whereas the strength of emotional valence representations in FPl correlated with emotion-driven benefits in task accuracy, i.e., increased affect-to-motor spillover. Collectively, these data suggest that FPl representations guide emotional context-appropriate behavior, while supporting the ability to disentangle emotional context from rule-guided cues.

### Conjunctive emotion-action representations in the basolateral amygdala

We also found evidence of conjunctive emotion-action representations in the basolateral amygdala. These results resonate with recent findings suggesting that amygdala neurons have mixed selectivity and represent contextual information. For instance, information about task sets, often observed in PFC neurons, is also decodable in single neurons in the non-human primate amygdala (Saez et al. 2015). Such abstract representations may provide important conceptual knowledge about when (and how) to respond to emotional events (Saez et al. 2015). Relatedly, amygdala intracranial event related potentials (iERPs) during an AGNG task represent the interaction of emotional valence (negative vs. neutral) and task rule (Go vs. No-Go) (Guex et al. 2020). Together, these results demonstrate a broader, more goal-oriented and contextually sensitive role for amygdala neurons than its more commonly emphasized emotional valence and arousal encoding functions (Pignatelli and Beyeler 2019).

In contrast to what we observed in FPl, in the amygdala, *incongruent* emotion-action trials were represented closer together in a high-multidimensional space compared to congruent trials. Moreover, when controlling for the emotion*action interaction regressor, aversive (fearful face) trials were represented as more *dissimilar* in the basolateral amygdala compared to positive (happy face) trials (in FPl, positive trials were represented as more dissimilar than aversive trials). Greater pattern dissimilarity in amygdala multivoxel patterns in response to negatively valenced stimuli has been observed in response to unpleasant odors (Jin et al. 2015), which may support early discrimination of aversive stimuli. The functional significance of representational distances—multivoxel pattern similarity vs. dissimilarity—is an ongoing area of study and varies by brain region (for instance, in hippocampus, pattern dissimilarity predicts better memory, in contrast to what is typically observed in neighboring brain regions such as perirhinal and parahippocampal cortices; (LaRocque et al. 2013; Copara et al. 2014)). It is possible that the observed dissimilarities for negative trials in the amygdala were due to competition of highly perceptually similar stimuli (i.e., negative and positive facial expressions of the same identity) and/or due to increased differentiation of specific face identities as they were repeated across the experiment; the design of the present study precluded disentangling these possibilities. Future work should further elucidate the temporal dynamics and behavioral significance of pattern dissimilarities in the amygdala and FPl.

### Clarifying the neuroanatomy of emotional valence integration into action goals

Extant theories of the hierarchical organization of cognitive control in LPFC propose that the frontopolar cortex (rostrolateral PFC) is a node through which internal states, putatively originating from vmPFC, arrive in LPFC (Badre and Nee 2018). Here, we examined this proposed pathway, and found converging evidence from functional connectivity and multivariate decoding analyses that a vmPFC region known to be densely interconnected with the amygdala and essential for emotion regulation, subgenual ACC (BA25), was the likely source of emotional signals arriving in FPl. (**Fig. 8**). First, the strength of multivariate decoding of emotion in BA25 significantly predicted the strength of emotion decoding in FPl. Secondly, BA25 and basolateral amygdala significantly coupled during negative emotional processing, consistent with known neuroanatomical projections (Ghashghaei, Hilgetag, and Barbas 2007; Barbas and Pandya 1989; Joyce and Barbas 2018). Further, the extent of functional coupling between BA25 and FPl during negative emotional processing predicted the strength of FPl’s multivariate decoding of emotional valence. These associations were not found in a neighboring vmPFC region, BA32, suggesting some degree of specificity of BA25-FPl interactions in this task. Collectively, these findings clarify the neuroanatomical pathway through which goal-based action may be potentiated or hindered by internal emotional signals arriving in LPFC, and how cognitive control is shaped by internal states. Moreover, these findings are consistent with recent work underscoring the importance of the amygdala ventrofugal pathway for the regulation of emotional action, and they deepen our understanding of the nodes in the (likely polysynaptic) circuitry linking amygdala and FPl function (Folloni et al. 2019; Bramson et al. 2020).

**Figure 8.**
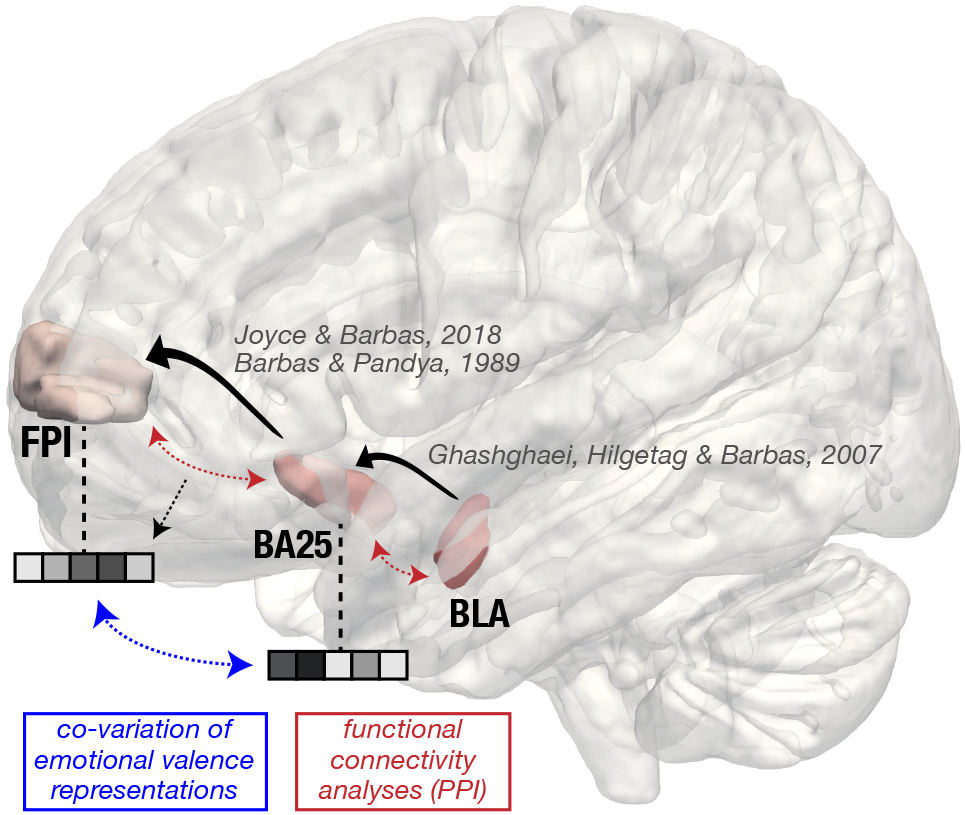
Summary of current results and extant neuroanatomical literature. Black arrows indicate neuroanatomical projections in the non-human primate, which are consistent with our findings from PPI and co-multivariate decoding analyses. The significant covariation of classifier evidence for emotional valence found between BA25 and the FPl is highlighted in blue, and PPI results are indicated in red.

### Implications for our understanding of LPFC organization, FPl function, and mental health

Emotional information comprises a critical source of contextual input that has been seldom considered in models of cognitive control in LPFC. This is despite the fact that LPFC lesions are known to predispose individuals to depression (Koenigs et al. 2008), LPFC TMS is an effective treatment for depression, particularly when its vmPFC projections are considered (Fox et al. 2012), and causal perturbations to LPFC via TMS impair automatic emotion regulation (Lapate et al. 2017).

In theories of the organization of cognitive control, FPl was once considered the top of a hierarchical gradient of goal abstraction, although recent work suggests that abstraction is distributed along various nodes in LPFC, and that mid-LPFC may be the apex of the LPFC cognitive control hierarchy (Derek Evan Nee and D’Esposito 2016; Derek E. Nee and D’Esposito 2017)—consistent with our finding that it held the strongest magnitude of action goal representations amongst the brain regions examined here. Nonetheless, it is clear that FPl’s integrative function cuts across stimulus domains in the service of complex behavior, as it is reliably engaged in conditions that involve relational processing and temporally abstract considerations, including alternative courses of action or counterfactuals (Badre and Nee 2018; Christoff and Gabrieli 2000; Boorman et al. 2009; Koch et al. 2018). Our study adds to this literature by highlighting an important instance of FPl’s integrative function, through which emotional states can shape long term goals and behavioral control with consequences for emotion regulation. Accordingly, recent work has shown that the extent to which FPl responds to the congruency of emotion and action goal (approach vs avoid) predicts stress responsivity and susceptibility to post-traumatic stress disorder (Kaldewaij et al. 2021, 2019).

### Limitations

The following limitations of the present study warrant caution and additional investigation. First, the AGNG task has a strong motor component that may obscure the unambiguous decoding of action goal (separate from motor action representations). While the pattern similarity analysis approach allowed us to explicitly control for motor confounds, the linear classifier analysis does not. Future work should use a task that matches movement demands across approach and avoidance conditions (e.g. (Bramson et al. 2020)). Second, action goals may engage different neural processes than other types of goals, such as those unfolding at longer time scales. Third, the associations reported here are inherently correlational. Future work using non-invasive brain stimulation will be critical for understanding the specific contributions of representations in FPl and LPFC for adaptive emotional behavior.

## Conclusion

This study unveils integrated emotion-action representations in FPl, which are linearly decodable and correlate with cognitive control performance during emotional processing. Collectively, these findings provide a deeper understanding of how emotions flexibly shape goal representations, a process that goes awry in psychopathy, as well as of how acute emotional states may disrupt goal-based emotion regulation in mood and anxiety disorders.

## Acknowledgments

The authors acknowledge the Wheeler Foundation, NIH grants MH63901 (MD), F32-MH113347 (RCL), F32-MH119796 (ICB), and NSF award BCS-0821855 (MD). The authors thank Jeanette Mumford, Jingyi Wang, Arielle Tambini, and Jacob Miller for helpful discussions, Lennart Verhagen for access to the PFC consensus atlas, and Audrey Phan, Jenna Martin and Jean Wu for assistance with data collection.

## References

Abraham, Alexandre, Fabian Pedregosa, Michael Eickenberg, Philippe Gervais, Andreas Mueller, Jean Kossaifi, Alexandre Gramfort, Bertrand Thirion, and Gaël Varoquaux. 2014. “Machine Learning for Neuroimaging with Scikit-Learn.” Frontiers in Neuroinformatics 8 (February): 14.

Badre, David, Apoorva Bhandari, Haley Keglovits, and Atsushi Kikumoto. 2021. “The Dimensionality of Neural Representations for Control.” Current Opinion in Behavioral Sciences 38 (April): 20–28.

Badre, David, and Mark D’Esposito. 2009. “Is the Rostro-Caudal Axis of the Frontal Lobe Hierarchical?” Nature Reviews. Neuroscience 10 (9): 659–69.

Badre, David, and Derek Evan Nee. 2018. “Frontal Cortex and the Hierarchical Control of Behavior.” Trends in Cognitive Sciences 22 (2): 170–88.

Ballard, Ian C., Anthony D. Wagner, and Samuel M. McClure. 2019. “Hippocampal Pattern Separation Supports Reinforcement Learning.” Nature Communications 10 (1): 1073.

Barak, Omri, Mattia Rigotti, and Stefano Fusi. 2013. “The Sparseness of Mixed Selectivity Neurons Controls the Generalization–Discrimination Trade-Off.” The Journal of Neuroscience: The Official Journal of the Society for Neuroscience 33 (9): 3844–56.

Barbas, H., and D. N. Pandya. 1989. “Architecture and Intrinsic Connections of the Prefrontal Cortex in the Rhesus Monkey.” The Journal of Comparative Neurology 286 (3): 353–75.

Baskin-Sommers, Arielle, Allison M. Stuppy-Sullivan, and Joshua W. Buckholtz. 2016. “Psychopathic Individuals Exhibit but Do Not Avoid Regret during Counterfactual Decision Making.” Proceedings of the National Academy of Sciences of the United States of America 113 (50): 14438–43.

Bates, Douglas, Martin Mächler, Ben Bolker, and Steve Walker. 2015. “Fitting Linear Mixed-Effects Models Using Lme4.” Journal of Statistical Software, Articles 67 (1): 1–48.

Bernardi, Silvia, Marcus K. Benna, Mattia Rigotti, Jérôme Munuera, Stefano Fusi, and C. Daniel Salzman. 2020. “The Geometry of Abstraction in the Hippocampus and Prefrontal Cortex.” Cell 183 (4): 954-967.e21.

Boorman, Erie D., Timothy E. J. Behrens, Mark W. Woolrich, and Matthew F. S. Rushworth. 2009. “How Green Is the Grass on the Other Side? Frontopolar Cortex and the Evidence in Favor of Alternative Courses of Action.” Neuron 62 (5): 733–43.

Bramson, Bob, Davide Folloni, Lennart Verhagen, Bart Hartogsveld, Rogier B. Mars, Ivan Toni, and Karin Roelofs. 2020. “Human Lateral Frontal Pole Contributes to Control over Emotional Approach– Avoidance Actions.” The Journal of Neuroscience: The Official Journal of the Society for Neuroscience 40 (14): 2925–34.

Christoff, Kalina, and John D. E. Gabrieli. 2000. “The Frontopolar Cortex and Human Cognition: Evidence for a Rostrocaudal Hierarchical Organization within the Human Prefrontal Cortex.” Psychobiology 28 (2): 168–86.

Cole, Michael W., Takuya Ito, and Todd S. Braver. 2016. “The Behavioral Relevance of Task Information in Human Prefrontal Cortex.” Cerebral Cortex 26 (6): 2497–2505.

Copara, Milagros S., Abdul S. Hassan, Colin T. Kyle, Laura A. Libby, Charan Ranganath, and Arne D. Ekstrom. 2014. “Complementary Roles of Human Hippocampal Subregions during Retrieval of Spatiotemporal Context.” The Journal of Neuroscience: The Official Journal of the Society for Neuroscience 34 (20): 6834–42.

Folloni, Davide, Jerome Sallet, Alexandre A. Khrapitchev, Nicola Sibson, Lennart Verhagen, and Rogier B. Mars. 2019. “Dichotomous Organization of Amygdala/Temporal-Prefrontal Bundles in Both Humans and Monkeys.” ELife 8 (November): e47175.

Fox, Michael D., Randy L. Buckner, Matthew P. White, Michael D. Greicius, and Alvaro Pascual-Leone. 2012. “Efficacy of Transcranial Magnetic Stimulation Targets for Depression Is Related to Intrinsic Functional Connectivity with the Subgenual Cingulate.” Biological Psychiatry 72 (7): 595–603.

Fusi, Stefano, Earl K. Miller, and Mattia Rigotti. 2016. “Why Neurons Mix: High Dimensionality for Higher Cognition.” Current Opinion in Neurobiology 37 (April): 66–74.

Ghashghaei, H. T., C. C. Hilgetag, and H. Barbas. 2007. “Sequence of Information Processing for Emotions Based on the Anatomic Dialogue between Prefrontal Cortex and Amygdala.” NeuroImage 34 (3): 905– 23.

Guex, Raphael, Constantino Méndez-Bértolo, Stephan Moratti, Bryan A. Strange, Laurent Spinelli, Ryan J. Murray, David Sander, Margitta Seeck, Patrik Vuilleumier, and Judith Domínguez-Borràs. 2020. “Temporal Dynamics of Amygdala Response to Emotion-and Action-Relevance.” Scientific Reports 10 (1): 1–16.

Jacobs, Bob, Matthew Schall, Melissa Prather, Elisa Kapler, Lori Driscoll, Serapio Baca, Jesse Jacobs, Kevin Ford, Marcy Wainwright, and Melinda Treml. 2001. “Regional Dendritic and Spine Variation in Human Cerebral Cortex: A Quantitative Golgi Study.” Cerebral Cortex 11 (6): 558–71.

Jenkinson, Mark, Christian F. Beckmann, Timothy E. J. Behrens, Mark W. Woolrich, and Stephen M. Smith. 2012. “FSL.” NeuroImage 62 (2): 782–90.

Jin, Jingwen, Christina Zelano, Jay A. Gottfried, and Aprajita Mohanty. 2015. “Human Amygdala Represents the Complete Spectrum of Subjective Valence.” The Journal of Neuroscience: The Official Journal of the Society for Neuroscience 35 (45): 15145–56.

Joyce, Mary Kate P., and Helen Barbas. 2018. “Cortical Connections Position Primate Area 25 as a Keystone for Interoception, Emotion, and Memory.” The Journal of Neuroscience: The Official Journal of the Society for Neuroscience 38 (7): 1677–98.

Kaldewaij, Reinoud, Saskia B. J. Koch, Mahur M. Hashemi, Wei Zhang, Floris Klumpers, and Karin Roelofs. 2021. “Anterior Prefrontal Brain Activity during Emotion Control Predicts Resilience to Post-Traumatic Stress Symptoms.” Nature Human Behaviour, February. https://doi.org/10.1038/s41562-021-01055-2.

Kaldewaij, Reinoud, Saskia B. J. Koch, Wei Zhang, Mahur M. Hashemi, Floris Klumpers, and Karin Roelofs. 2019. “Frontal Control Over Automatic Emotional Action Tendencies Predicts Acute Stress Responsivity.” Biological Psychiatry: Cognitive Neuroscience and Neuroimaging 4 (11): 975–83.

Kamali, Arash, Haris I. Sair, Ari M. Blitz, Roy F. Riascos, Saeedeh Mirbagheri, Zafer Keser, and Khader M. Hasan. 2016. “Revealing the Ventral Amygdalofugal Pathway of the Human Limbic System Using High Spatial Resolution Diffusion Tensor Tractography.” Brain Structure & Function 221 (7): 3561–69.

Kikumoto, Atsushi, and Ulrich Mayr. 2020. “Conjunctive Representations That Integrate Stimuli, Responses, and Rules Are Critical for Action Selection.” Proceedings of the National Academy of Sciences of the United States of America 117 (19): 10603–8.

Kim, Maxwell L. Elliott, Annchen R. Knodt, and Ahmad R. Hariri. 2020. “A Connectome-Wide Functional Signature of Trait Anger.” BioRxiv. https://doi.org/10.1101/2020.10.14.338863.

Koch, Saskia B. J., Rogier B. Mars, Ivan Toni, and Karin Roelofs. 2018. “Emotional Control, Reappraised.” Neuroscience and Biobehavioral Reviews 95 (December): 528–34.

Koenigs, Michael, Edward D. Huey, Matthew Calamia, Vanessa Raymont, Daniel Tranel, and Jordan Grafman. 2008. “Distinct Regions of Prefrontal Cortex Mediate Resistance and Vulnerability to Depression.” The Journal of Neuroscience: The Official Journal of the Society for Neuroscience 28 (47): 12341– 48.

Kriegeskorte, Nikolaus, and Rogier A. Kievit. 2013. “Representational Geometry: Integrating Cognition, Computation, and the Brain.” Trends in Cognitive Sciences 17 (8): 401–12.

Lapate, Regina C., Jason Samaha, Bas Rokers, Hamdi Hamzah, Bradley R. Postle, and Richard J. Davidson. 2017. “Inhibition of Lateral Prefrontal Cortex Produces Emotionally Biased First Impressions: A Transcranial Magnetic Stimulation and Electroencephalography Study.” Psychological Science 28 (7): 942–53.

LaRocque, Karen F., Mary E. Smith, Valerie A. Carr, Nathan Witthoft, Kalanit Grill-Spector, and Anthony D. Wagner. 2013. “Global Similarity and Pattern Separation in the Human Medial Temporal Lobe Predict Subsequent Memory.” The Journal of Neuroscience: The Official Journal of the Society for Neuroscience 33 (13): 5466–74.

Medalla, Maria, and Helen Barbas. 2010. “Anterior Cingulate Synapses in Prefrontal Areas 10 and 46 Suggest Differential Influence in Cognitive Control.” The Journal of Neuroscience: The Official Journal of the Society for Neuroscience 30 (48): 16068–81.

Mumford, Jeanette A., Benjamin O. Turner, F. Gregory Ashby, and Russell A. Poldrack. 2012. “Deconvolving BOLD Activation in Event-Related Designs for Multivoxel Pattern Classification Analyses.” NeuroImage 59 (3): 2636–43.

Nee, Derek E., and Mark D’Esposito. 2017. “Causal Evidence for Lateral Prefrontal Cortex Dynamics Supporting Cognitive Control.” ELife 6 (September): e28040.

Nee, Derek Evan, and Mark D’Esposito. 2016. “The Hierarchical Organization of the Lateral Prefrontal Cortex.” ELife 5 (March). https://doi.org/10.7554/eLife.12112.

Neubert, Franz-Xaver, Rogier B. Mars, Adam G. Thomas, Jerome Sallet, and Matthew F. S. Rushworth. 2014. “Comparison of Human Ventral Frontal Cortex Areas for Cognitive Control and Language with Areas in Monkey Frontal Cortex.” Neuron 81 (3): 700–713.

O’Reilly Jill X., Mark W. Woolrich, Timothy E. J. Behrens, Stephen M. Smith, and Heidi Johansen-Berg. 2012. “Tools of the Trade: Psychophysiological Interactions and Functional Connectivity.” Social Cognitive and Affective Neuroscience 7 (5): 604–9.

Pignatelli, Michele, and Anna Beyeler. 2019. “Valence Coding in Amygdala Circuits.” Current Opinion in Behavioral Sciences 26 (April): 97–106.

Reuter, Martin, Nicholas J. Schmansky, H. Diana Rosas, and Bruce Fischl. 2012. “Within-Subject Template Estimation for Unbiased Longitudinal Image Analysis.” NeuroImage 61 (4): 1402–18.

Saez, A., M. Rigotti, S. Ostojic, S. Fusi, and C. D. Salzman. 2015. “Abstract Context Representations in Primate Amygdala and Prefrontal Cortex.” Neuron 87 (4): 869–81.

Sallet, Jérôme, Rogier B. Mars, Maryann P. Noonan, Franz-Xaver Neubert, Saad Jbabdi Jill, X. O’Reilly, Nicola Filippini, Adam G. Thomas, and Matthew F. Rushworth. 2013. “The Organization of Dorsal Frontal Cortex in Humans and Macaques.” Journal of Neuroscience 33 (30): 12255–74.

Smith, Stephen M., Mark Jenkinson, Mark W. Woolrich, Christian F. Beckmann, Timothy E. J. Behrens, Heidi Johansen-Berg, Peter R. Bannister, et al. 2004. “Advances in Functional and Structural MR Image Analysis and Implementation as FSL.” NeuroImage 23 Suppl 1: S208–19.

Tye, Kay M. 2018. “Neural Circuit Motifs in Valence Processing.” Neuron 100 (2): 436–52.

Tyszka, J. Michael, and Wolfgang M. Pauli. 2016. “In Vivo Delineation of Subdivisions of the Human Amygdaloid Complex in a High-Resolution Group Template.” Human Brain Mapping 37 (11): 3979– 98.

Verhagen, Lennart. 2018. “Prefrontal Consensus Atlas (Oxford).” http://lennartverhagen.com/;lennart.verhagen@donders.ru.nl.

Volman, Inge, Karin Roelofs, Saskia Koch, Lennart Verhagen, and Ivan Toni. 2011. “Anterior Prefrontal Cortex Inhibition Impairs Control over Social Emotional Actions.” Current Biology: CB 21 (20): 1766– 70.

Walther, Alexander, Hamed Nili, Naveed Ejaz, Arjen Alink, Nikolaus Kriegeskorte, and Jörn Diedrichsen. 2016. “Reliability of Dissimilarity Measures for Multi-Voxel Pattern Analysis.” NeuroImage 137 (August): 188–200.

Waskom, Michael L., Dharshan Kumaran, Alan M. Gordon, Jesse Rissman, and Anthony D. Wagner. 2014. “Frontoparietal Representations of Task Context Support the Flexible Control of Goal-Directed Cognition.” The Journal of Neuroscience: The Official Journal of the Society for Neuroscience 34 (32): 10743–55.

